# *Asip* (Agouti-signaling protein) aggression gene regulate auditory processing genes in mice

**DOI:** 10.1101/2020.06.10.141325

**Authors:** Alexandra Mickael, Pavel Klimovich, Patrick Henckel, Norwin Kubick, Michel E Mickael

## Abstract

Covid-19 strategy of lockdown has affected the lives of millions. The strict actions to enclose the epidemic have exposed many households to inner tensions. Domestic violence has been reported to increase during the lockdown. However, the reasons for this phenomenon have not been thoroughly investigated. Melanocortin GPCRs family contribution to aggression is well documented. ASIP (nonagouti) gene plays a vital role in regulating the melanocortin GPCRs family function, and it is responsible for regulating aggression in mice. We conducted a selection analysis of ASIP. We found that it negatively purified from Shark to humans. In order to better asses the effect of this gene in mammals, we performed RNA-seq analysis of a knockout of an ASIP crisper-cas mouse model. We found that ASIP KO in mice upregulates several genes controlling auditory function, including *Phox2b*, *Mpk13*, *Fat2, Neurod2, Slc18a3, Gon4l Gbx2, Slc6a3(Dat1) Aldh1a7 Tyrp1 and Lbx1*. Interestingly, we found that *Slc6a5*, and *Lamp5* as well as *IL33*, which are associated with startle disease, are also upregulated in response to knocking out ASIP. These findings are indicative of a direct autoimmune effect between aggression-associated genes and startle disease. Furthermore, in order to validate the link between aggression and auditory inputs processing. We conducted psychological tests of persons who experienced lockdown. We found that aggression has risen by 16 % during the lockdown. Furthermore, 3% of the subjects interviewed reported a change in their hearing abilities. Our data shed light on the importance of the auditory input in aggression and open perceptions to interpret how hearing and aggression interact at the molecular neural circuit level.

## Introduction

The link between social lockdown and aggression is well established. Aggression can be categorized as a beneficial evolutionary trait as it might indicate survival when individuals are competing for resources. Conversely, aggressiveness might also impede social cohesion. Social lockdown can lead to psychological problems, including anxiety, depression, antisocial, and violence-related behaviors. For example, orca confinement in closed places exhibits aggressive behaviors [1]. In humans, release from mandatory confinement indoors was correlated with decreases in both verbal and physical aggression[2, 3]. During Covid-19 lockdown, 45.3% of participants of a study investigating the effect of lockdown reported worsened sleep, increased levels of irritation, anger, and aggression compared to pre-pandemic times[3]. Furthermore, 5% of all participants reported experiencing interpersonal violence (IPV)[3]. However, the factors causing aggression are not yet well investigated.

Melanocortin system plays an important role in regulating aggression. It is widely documented that the peptide hormone pro-opiomelanocortin (POMC) acts as a precursor for various (neuro)peptides including α- melanocyte-stimulating hormone (α-MSH), adreno-corticotrophin (ACTH) and β-endorphin. The function of melanocortins is regulated by the activation of a family of melanocortin receptor subtypes. α- MSH binds to MCRs in the brain, where it regulates social behavior, appetite and stress physiology. α- MSH acts as a neurotransmitter in the brain where it can modulate behavior mainly via central MC3R and MC4R in a variety of ways including regulating the dopaminergic reward system. Melanocortin-5 receptor deficiency reduces a pheromone signal for aggression in male mice[4]. Interestingly, agouti signaling peptide (*Asip*) and agouti-related peptide (*Agrp*) have diverse functional roles in feeding, pigmentation and background adaptation mechanisms. Interestingly, ASIP serves as anof the MC1R and the MC4R receptors. It seems melancortin system diverged from adenosine receptors around the time of divergence of *Hydra vulgaris* [5]. Serotonin receptors that are known to play a distinctive role in aggression seemed to have evolved as they have been found in *Trichoplax adhereans* [5]. While dopamine receptors diverged afterwards around the time of emergence of *Ciona intestinalis* [5]. However, the inter-regulation mechanism between these four families is still not clear.

Hearing as a sensory modality in the context of aggressive behavior has been shown to play a major role in controlling behavior [6]. Precise integration and processing of sensory inputs are crucial to evoke a suitable behavioral response [7]. In crickets, aggression songs are associated with cricket fights[8]. Mice lacking *Asic3* show reduced anxiety◻like behavior on the elevated plus maze and reduced aggression [9]. *Asic3* was also shown to affect hearing[10]. These reports indicate that aggression and hearing are possibly interlinked. Previous reports have shown various sensory modalities in mediating of aggressive behavior in *Drosophila melanogaster* including olfactory, gustatory, as well as visual neural networks[6]. Furthermore it was found that, neuronal silencing and targeted knockdown of hearing genes such as d trpl (transient receptor potential-like) and the Ca2+ signaling-related genes Arr2 and inaD in the fly’s auditory organ elicit abnormal aggression[6]. These observation indicates that hearing could regulating aggression behavior. However if aggression controls hearing is not yet known. Interestingly, Melanocytes present in the cochlea have an essential function in inner ear physiology. They protect against various types of hearing loss, including age-related hearing loss (ARHL) and noise-related hearing loss (NIHL), by means of calcium buffering, heavy metal scavenging, and antioxidant activities [11]. However if aggressiveness is directly affecting hearing abilities is still not known. Furthermore there has no been earlier reports of Asip investigations that have shown a direct link between ASIP and hearing.

In this study, we have investigated the role of *Asip* in linking aggression and the hearing system. In order to confirm that our mice model study would be representative of aggression hearing link, we conducted an evolutionary study that revealed that ASIP is negatively selected between mice and humans. We analyzed an RNA-seq data in which ASIP was knocked out from Japanese wild-derived MSM/Ms strain using CRISPR/Cas9 genome editing. We found that numerous hearing associated genes were upregulated in the KO mouse model including those linked to startle disease. In order to validate our results we conducted behavioral tests, to investigate whether the rise in aggression during the covid-19 period has affected hearing patterns, we found that there 5 % individuals interviewed experience a change in their hearing abilities while engaging in arguments during lockdown.

## Methods

### Alignment and phylogenetic analysis

Phylogenetic investigation was done in three stages. First, ASIP family amino acid sequences were aligned using MAFFT via the iterative refinement method (FFT-NS-i). Next, we employed ProtTest to conclude the best amino acid replacement model [12]. ProtTest results based on the Akaike information criterion (AIC) suggested that the best substitution model is LG+I+G+F., LG is the substitution model supplemented by a fraction of invariable amino acids (‘+I’) with each site assigned a probability of belonging to given rate categories (‘+G’) and observed amino acid frequencies (‘+F’). The third stage involved employing the protein alignment and the resulting substitution model, in applying two different phylogenetic methods to construct the tree, namely, (1) maximum likelihood and (2) Bayesian inference. We performed the maximum likelihood analysis using PHYML[13] implemented in Seaview with 5 random starting trees[14]. We applied Bayesian inference analysis using MrBayes where we implemented a Markov chain Monte Carlo analysis with 1000,000 generations to approximate the posterior probability and a standard deviation of split frequencies <0.01 to indicate convergence as previously described. We used the coding DNA alignment and our final tree to investigate the ratio of non-synonymous (dN) to synonymous (dS) amino acid substitutions using the PAML program. Likelihood ratio tests (LRT) were constructed to compare the p-values of χ2 square tests for selective pressure models against neutral models. One level of analysis was investigated. This level calculates the global ω for the tree using the one-ratio model M0[15] where ω = dN/dS, with trees under purifying selection (0<dN/dS<1) as well as trees under neutral evolution (dN/dS=1), while the positive selection model evolved under (dN/dS >1).

### Analysis of RNA-seq Data

The overall design of the RNA experiment was as follows: The mid brain section was isolated from Japanese wild-derived MSM/Ms Japanese mice (2 control and 3 KO)[16]. Total RNA was purified from dissected midbrain using Trizol (Thermo Fisher Scientific). Purified total RNA was amplified and labeled with Cy3 using Low-Input QuickAmp Labeling Kit (Agilent Technologies). Cy3-labeled RNAs were hybridized to SurePrint G3 Mouse Gene Expression v2 8×60K Microarray Kit (Agilent Technologies) at 65 °C for 17h. The scanned images were processed with Feature Extraction software (Agilent Technologies) to extract signal intensities of each probe. The extracted signal data were imported into the Gene Spring GX 13.1.1 software (Agilent Technologies) and normalized using the default settings. RNA-seq analysis was then performed in R using Limma [17]. Briefly, we employed the Limma RNA-seq differential gene expression method to compute the non-parametric approximations of mean–variance relationships. This allowed us to calculate the weights for a linear model analysis of log-transformed counts in conjunction with the empirical Bayes shrinkage of variance parameters. Differential expression analysis was performed to determine the differences in gene expression between +LPS cells and non-treated samples by fitting a linear model to compute the variability in the data with LMFIT [72,73]. Pathway enrichment was done using the library FGSEA[18]. The network between chosen genes was calculated using the GLASSO module utilizing the webserver GeNeCK [19] with default settings.

### Human Proteome Atlas analysis

Data were downloaded from http://www.proteinatlas.org [20]. For a protein to be a candidate biomarker it should be medium or high expressed in normal brain. We arbitrarily set our selection criteria for candidate genes that were found to be upregulated in the RNA-seq study.

### Sample

The present study included 25 participants who reported no prior aggression, or depression diagnosis. The sample size was determined by the G power analysis. The participants reported age ranged between 27 to 65 years. The education level of participants varied from uneducated to master’s degree. Informed written consent was taken from the participants after explaining them the purpose of research. **Exclusion criteria;** Participants with some serious general medical condition were also not selected.

### Aggression measurement

Aggression was measured Buss-Perry Aggression Questionnaire[21]. This scale consists of four subscales of physical aggression verbal aggression, anger and hostility The participants were asked to evaluate each item on a 5-point Likert scale ranging from 1 (not true) to 5 (true). The consistency for each category was confirmed in with Cronbach alphas ranging from .70 to .78[22]. In this study, the alphas for the subscales were .75 for physical aggression, .77 for verbal aggression, .76 for anger, and .73 for hostility, respectively. The descriptive scores of the four subscales were considered by averaging the item scores.

### Bergen Social Relationship

We examined whether aggression behavior could be caused by social stress. We employed the Bergen Social Relationship Scale (BSRS) which measures the interpersonal relationship problems[23]. It is a six element self-report quantity. The answers were on four points scale using the notions of “describe me well (3)” to “do not describe me at all”. The scores system was divided from 0 to 18, where a higher score signifies higher interpersonal conflicts. Cronbach’s alpha for the BSRS was reported to be 0.76. The test-retest correlation was reported to be 0.75. The construct validity of the BSRS was ranged from 0.40 to 0.32, all statistically significant at P < 0.001.

## Results

Our results indicate that aggression-inducing gene the conserved aggression genes *Asip* is responsible for down regulating several genes responsible for hearing and acoustic processing in mice. Our results also shows that this gene is strongly conserved between mice and humans. When we analyzed RNA-seq for Asip ko mouse model we found that several genes controlling hearing were upregulated in the KO samples. Specifically, *Phox2b*, *Mpk13*, *Fat2, Neurod2, Slc18a3, Gon4l Gbx2, Slc6a3 Aldh1a7 Tyrp1* and *Lbx1* genes were down regulated in *Asip (ko)*. Interestingly, we found that *Slc6a5*, and *Lamp5* as well as *IL33*, which are associated with startle disease, are also upregulated in response to knocking out ASIP. These results imply that Asip is playing a fundamental role in startle disease and that the startle disease pathology is connected to the patient’s response to aggression. We found the link between hearing abilities and aggression mirrored in human samples where people whose experienced aggression behavior lockdown, also reported hearing abilities change. Our results shed more light on the link between aggression and processing acoustic signals in humans.

### Evolution results

Agouti evolution was under negative selection. Our results extend what has been reported by Saeed et al. We were able to locate human *Asip* gene homologs in chimps, orangutan, mice, and chicken (Figure x). We found two homologs of the gene in zebrafish as well as in elephant shark (FAA00702.1, and FAA00750.1). Interestingly, we were only able to find a single copy of the gene in the reed fish Erpetoichthys calabaricus (XP_028660269.1), suggesting that Reed fish lost one homolog. We were not able to locate any Asip genes in Lampreys. However, we were able to locate several melanocortin receptors (e.g. XP_032827023.1, serotonin, XP_032806630.1, adenosine (XP_032823932.1) and dopamine receptors XP_032806557.1. Suggesting that lampreys have evolved its unique pathways for regulation of aggression. We were not able to locate *Asip* in Hagfish Hyperotreti, Ciona intestinalis, Hydra Vulgaris, Drosophila melanogaster, Trichoplax adhaerens or Caenorhabditis elegans. These observations indicate that Asip first emerged during the divergence of Gnathostomata and Hyperoartia (lampreys). Identity and similarity analysis indicated that *Asip* is highly conserved. For example, the similarity between humans chimp for *Asip* was 97.7 %, while with the mouse, it was (75.5%) (Figure). Furthermore, w ds/dn showed a value of 0.3, confirming that the genes evolved under negative selection (Figure x).

### Knocking out aggression gene results in increasing acoustic processing genes

We analyzed the GEO dataset GSE84840. In this dataset, CRISPR/Cas9-mediated genome editing in wild-derived mice was performed to generate tamed wild-derived strains by mutation of the a (nonagouti) gene. These tamed mice show non aggressive behavior when tested through the () test. We found genes related to hearing and acoustic signals processing upregulated in the ko mice. We also found that serotonin level was directly down regulated by knocking out agouti gene (figure 1). Other aggressiveness related genes that were down regulated include (). However interestingly we found serveal genes that are associated with aggressive behavior have not been affected such as cfos, and…. Notably, we did not see any change in the inflammatory pathway in this particular model. However, using the GEO data set () that uses (), we found that CD4 pathway was affected (figure 2) as well as () patwhay. (figure 2). (Figure 3=IHC)

**Figure 1.**
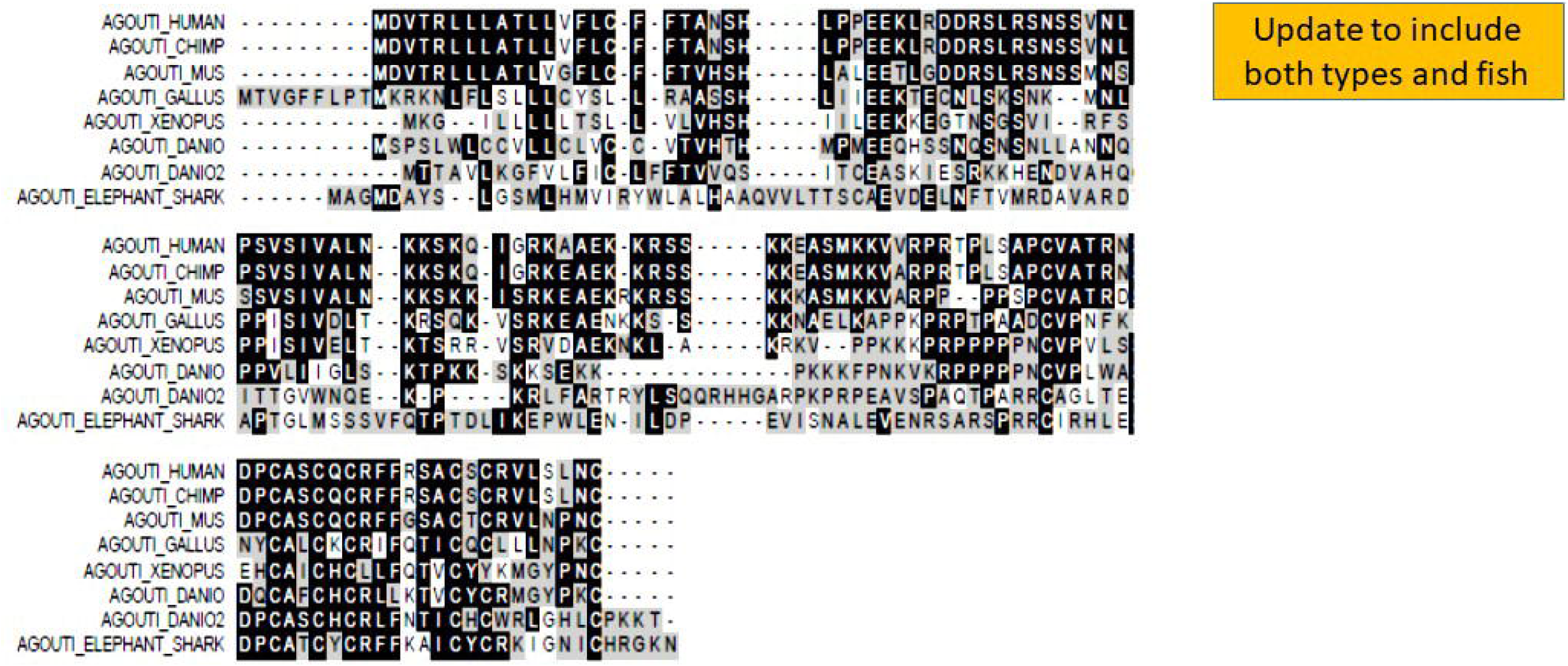
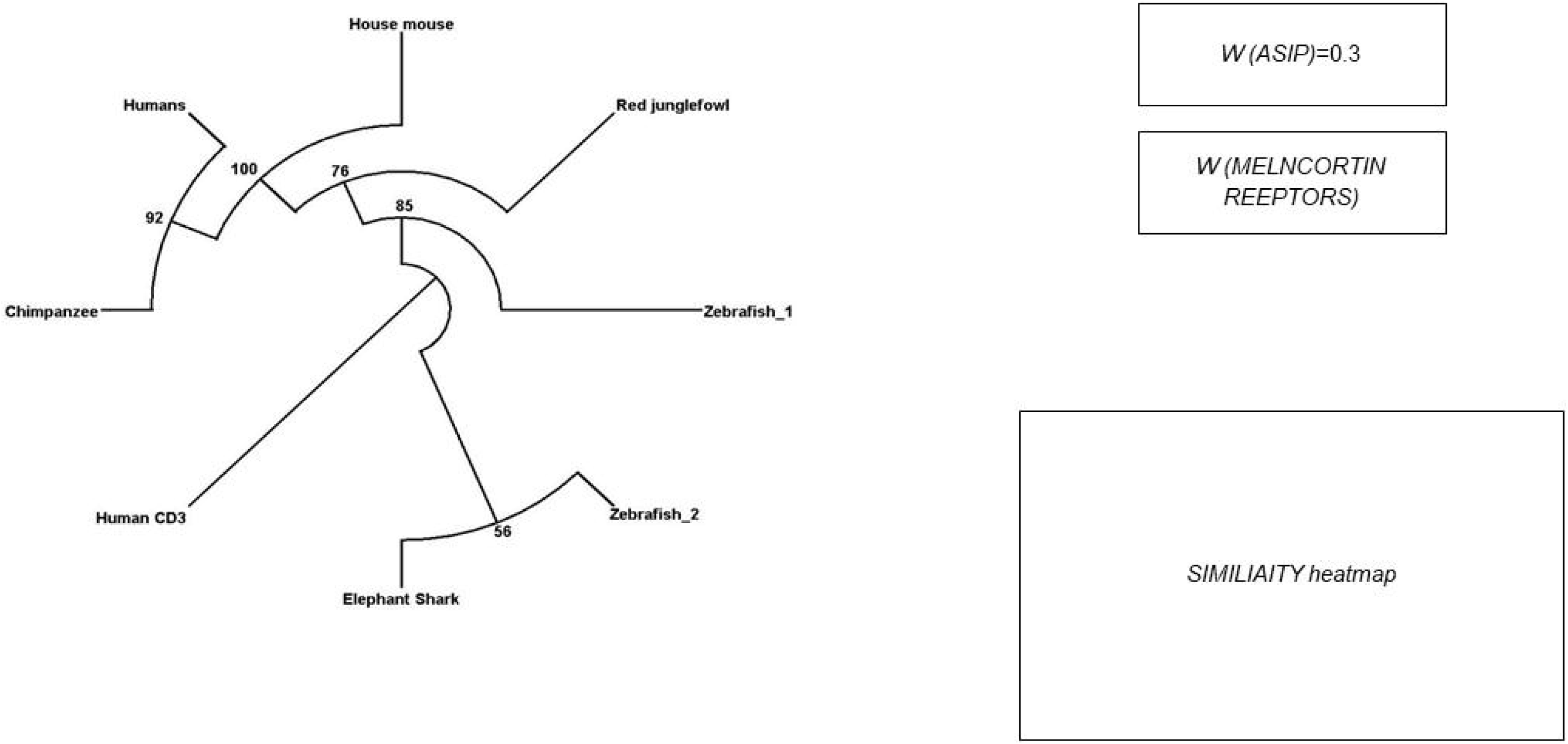
alignment high conservation level of Asip. **b)** Phylogenetic tree of nonagouti homologous proteins. Cartilaginous fishes such as Elephant shark were the first organisms to show nonagouti homologs. Zebra fish shows duplication events (2R). We are also able to identify nonagouti, in birds and rodents and primates. Human CD3 was used as a root. Phylogenetic bootstrap values are indicated adjacent to each node. **C)** Agouti gene is under negative selection (w 0.3) <1. (ADD memebrane annotation, consensus sequence). Figure x. **b)** Phylogenetic tree of nonagouti homologous proteins. Cartilaginous fishes such as Elephant shark were the first organisms to show nonagouti homologs. Zebra fish shows duplication events (2R). We are also able to identify nonagouti, in birds and rodents and primates. Human CD3 was used as a root. Phylogenetic bootstrap values are indicated adjacent to each node. **C)** Agouti gene is under negative selection (w 0.3) <1. (ADD memebrane annotation, consensus sequence) 1d) high similarity between

**Figure 2.**
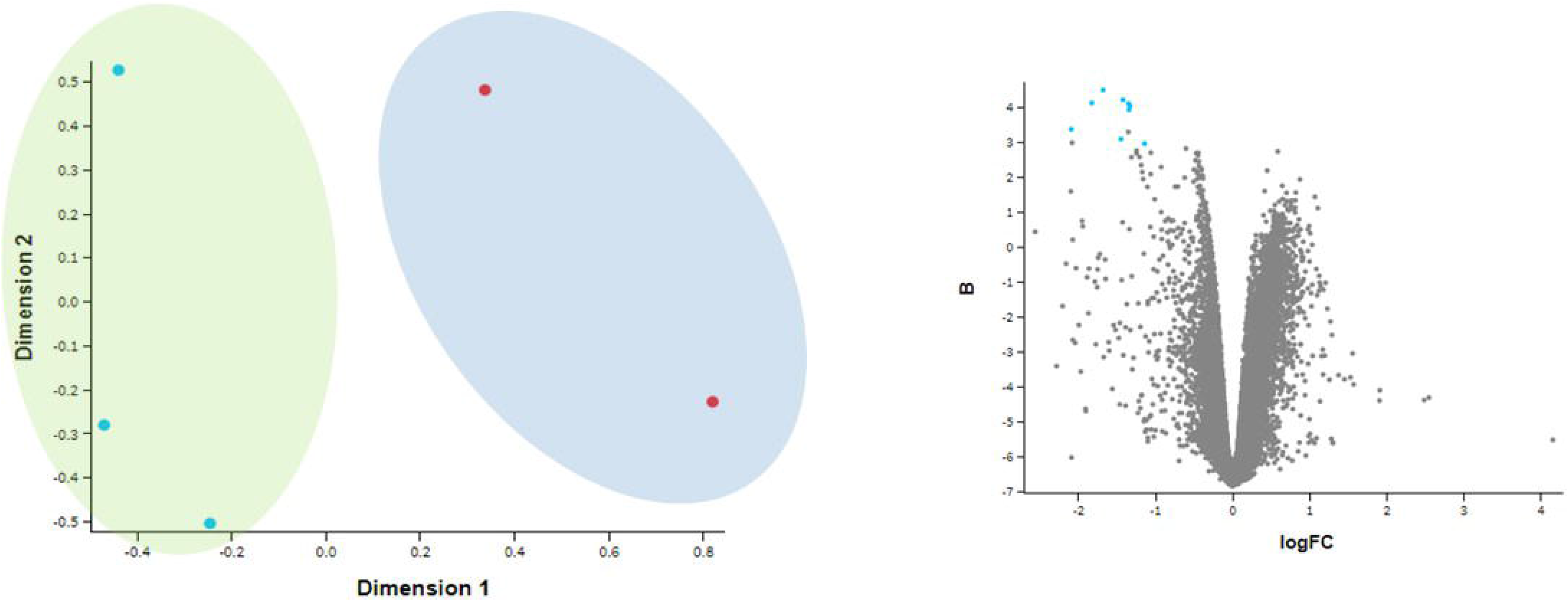
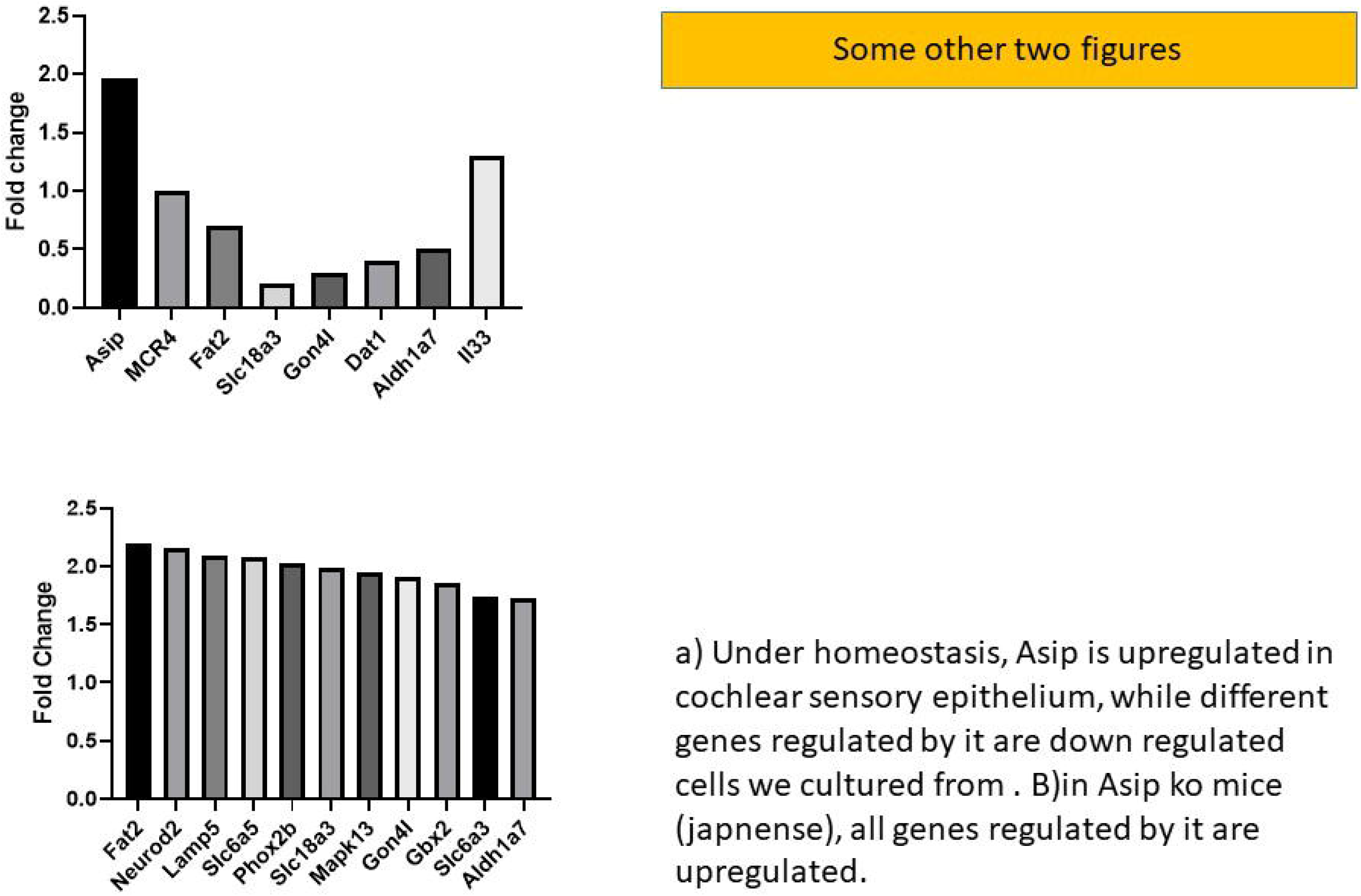
alignment high conservation level of Asip. **b)** Phylogenetic tree of nonagouti homologous proteins. Cartilaginous fishes such as Elephant shark were the first organisms to show nonagouti homologs. Zebra fish shows duplication events (2R). We are also able to identify nonagouti, in birds and rodents and primates. Human CD3 was used as a root. Phylogenetic bootstrap values are indicated adjacent to each node. **C)** Agouti gene is under negative selection (w 0.3) <1. (ADD memebrane annotation, consensus sequence). Figure x. **b)** Phylogenetic tree of nonagouti homologous proteins. Cartilaginous fishes such as Elephant shark were the first organisms to show nonagouti homologs. Zebra fish shows duplication events (2R). We are also able to identify nonagouti, in birds and rodents and primates. Human CD3 was used as a root. Phylogenetic bootstrap values are indicated adjacent to each node. **C)** Agouti gene is under negative selection (w 0.3) <1. (ADD memebrane annotation, consensus sequence) 1d) high similarity between

**Figure 3.**
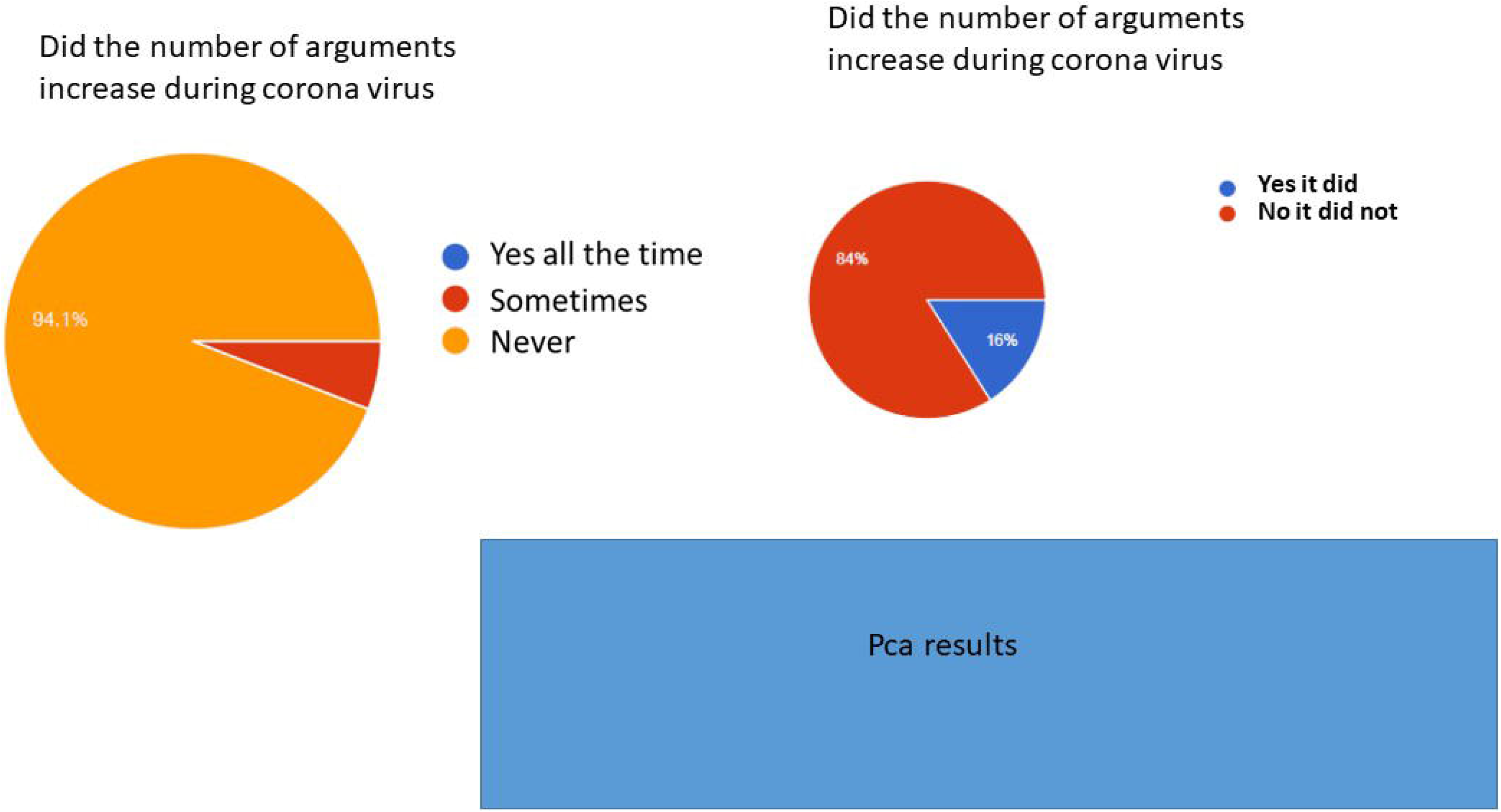
alignment high conservation level of Asip. **b)** Phylogenetic tree of nonagouti homologous proteins. Cartilaginous fishes such as Elephant shark were the first organisms to show nonagouti homologs. Zebra fish shows duplication events (2R). We are also able to identify nonagouti, in birds and rodents and primates. Human CD3 was used as a root. Phylogenetic bootstrap values are indicated adjacent to each node. C) Agouti gene is under negative selection (w 0.3) <1. (ADD memebrane annotation, consensus sequence). Figure x. **b)** Phylogenetic tree of nonagouti homologous proteins. Cartilaginous fishes such as Elephant shark were the first organisms to show nonagouti homologs. Zebra fish shows duplication events (2R). We are also able to identify nonagouti, in birds and rodents and primates. Human CD3 was used as a root. Phylogenetic bootstrap values are indicated adjacent to each node. **C)** Agouti gene is under negative selection (w 0.3) <1. (ADD memebrane annotation, consensus sequence) 1d) high similarity between

### Asip is expressed in the cochlear sensory epithelia

To investigate the role of Asip in the auditory sensory mechanisms, we analyzed the public Microarray of cochlear sensory epithelia versus embryonic stem cells. We found that Asip is upregulated in cochlear sensory epithelia but not in embryonic stem cells (fold change 1.92). Interestingly MCR4 was not upregulated (1.07) indicative of Asip a regulatory role of melanocortin receptors under homeostasis. Fat2, *Slc18a3, Gon4l, Dat1, Aldh1a7* were also not upregulated confirming our hypothesis of a regulatory effect of Asip on these hearing related genes.

### Asip is upregulated LPS treated cochlear sensory epithelia

To investigate the role of Asip in the cochlear sensory epithelia, we analyzed the public Microarray of cochlear sensory epithelia treated with LPL. We found that Asip is upregulated in cochlear sensory epithelia but not in embryonic stem cells (fold change 1.92). Interestingly IL 33 known to perform regulation of inflammatory pathway was not upregulated. Interestingly MCR4 was not upregulated (1.07) indicative of Asip a regulatory role of melanocortin receptors under homeostasis. Fat2, *Slc18a3, Gon4l, Dat1, Aldh1a7* were also not upregulated confirming our hypothesis of a regulatory effect of Asip on these hearing related genes.

#### Expression of proteOnaragraph (questionnaire)

**Table 1.** Results of the questionnaire suggests high hostility caused by lockdown.

| type | results | Meaning |
| --- | --- | --- |
| anger | 3.3 | Moderate (0-5) ( 5 is highest) |
| Physical aggression | 3.3 | Moderate (0-5) ( 5 is highest) |
| Hostile | 3.6 | High (0-5) ( 5 is highest) |
| Verbal aggression | 3.5 | High (0-5) ( 5 is highest) |
| Stress | 2.56 | Low (0-4) (0 is highest) |

## Discussion

Our studies suggest a direct link between acoustic processing and aggression. We have investigated the relationship between aggression induced by lockdown and hearing. We found that 3% of individuals who answered the questionnaire reported some difference in their hearing ability. This includes both and negative abilities. We found these results reflected in the molecular pathway of melanocortin receptors 1 and 4, where their knocking out their negative agonist; *Asip* resulted in upregulating various hearing associated genes such as *Fat2* among others. We also noticed that of the genes associated with hearing affected by knocking out of Asip is *Slc6a5* and *IL33*, which showed upregulation; both these genes were associated with startle disease. We investigated Asip further and we found that it is expressed in cochlear sensory epithelial cells. From an evolutionary point of view, Asip is more recent than melanocortin receptors. Taken together, our results suggest that *Asip* divergence represent the evolution of new mechanism linking hearing and aggression in higher animals.

### Evolution of the melanocortin system

Our analysis has shown that the *Asip* gene diverged around 600 MYA ago. *Ingo et Al*, (PMID: 21406593) have shown that Agouti exist in teleost. We have found it in elasmobranchs, proving it was more ancient than previously thought. The endocrine related genes that play an important role in aggression include, serotonin, with low levels of serotonin associated with violent behavior and suicidal thoughts. Serotonin perform its role in aggression through a network of genes including *Maoa* and *Maob*, (which play a role in the metabolism of biogenic amines), *Slc6a4*, (which acts as Serotonin transporter) and Tryptophan hydroxylase enzyme (*Tph1*) (which catalyzes the rate-limiting step in the synthesis of serotonin). It has been shown that polymorphism of metabolic enzymes, carrier proteins, and receptors on the serotonergic system are associated with an increased aggressive behavior pattern. Interestingly, serotonin receptors diverged during the time of *Trichoplax adhaeren*s. In human studies, a positive relationship between aggressiveness 3-methoxy-4-hydroxyphenylglycol (MHPG) (norepherinrine product) and has been established. Noradrenaline transporter (SLC6A2), −3081T allele is more dominate in ADHD in Americans of European descent, thus proving a questionable link with aggression. Interestingly, β-type noradrenergic receptor blocker have been shown to reduce aggressive behavior in some but not all patients tested, suggesting that aggression is controlled by a host of gene networks. Adrenergic receptors also evolved during the time of evolution of *Trichoplax adhaerens*. It seems the melanocortin system diverged from adenosine receptors during the divergence of hydra Vulgaris (PMID: 27614546). Conversely, arginine vasopressin levels are positively correlated with life history of aggression have evolved ruing the emergence of *Ciona intestinalis.* Another important endocrine mechanism is the dopamine reward system, for example, the gene DBH is a key enzyme in the synthesis of norepinephrine which is associated with conduct disorder. However, dopamine receptors first appeared during the emergence of *Ciona intestinalis*. Acetylcholine receptors also implicated in aggression behavior diverged during *Danio rerio* emergence. We could not find *Asip* in *Ciona intestinalis*. We have noticed that knocking out *Asip* increased acetylcholine receptor *Slc18a3* (2.1 fold increase). Interestingly, knocking out Asip did not affect dopaminergic or serotonergic receptors expression. Since its emergence *Asip* has been subjected to a tight negative selection (w=0.23). This indicates that from a chronological point of view, serotonin, Adrenergic, are responsible for regulating the basic mechanism controlling behavior in lower organisms, melanocortin and dopamine emerged as the need for a better control of aggression appeared in Hydra and *Ciona*, while acetylcholine receptors play a role in higher animals. Finally, Asip emerged to play a role of regulator in higher invertebrates and vertebrates.

#### *Asip* is controlling melanocortin role in hearing

Effect on *Asip* on genes associated with auditory signal processing. *Asip* is downregulating genes responsible for sound processing and regulation. *Phox2b*, *Mpk13*, *Fat2, Neurod2, Slc18a3, Gon4l Gbx2, Slc6a3(Dat1) Aldh1a7 Tyrp1 and Lbx1.* PHOX2B is expressed in the brain stem, mutation in this gene have been associated with brainstem dysfunction and brainstem auditory evoked potentials in 20% of the Congenital central hypoventilation syndrome (CCHS) patients investigated (25886294). PHOX2B is involved in the development of several major noradrenergic neuron populations, including the locus coeruleus the pons of the brainstem which is known to play a major role in aggression behavior (27083854). Fat2 plays an important role in hair cell regeneration in Zerbra fish (24706895). It was demonstrated that the lateral and basolateral amygdala nuclei fail to form in neuroD2 null mice and neuroD2 heterozygotes have fewer neurons in this region. NeuroD2 heterozygous mice show profound deficits in emotional learning as assessed by fear conditioning (16203979). Human NeuroD2can induce neurogenic differentiation in non-neuronal cells in Xenopus embryos, and is thought to play a role in the determination and maintenance of neuronal cell fates. Gbx2 Is Required for the Morphogenesis of the Mouse Inner Ear. In particular, absence of the endolymphatic duct and swelling of the membranous labyrinth are common features in Gbx2-/- inner ears. More severe mutant phenotypes include absence of the anterior and posterior semicircular canals, and a malformed saccule and cochlear duct(15829521). In slc6a3:EGFP larvae, it was found that EGFP-positive dopaminergic fibers were located within the supporting cell layer and not within the hair cell layer. It was demonstrated that dopamine receptors are present in sensory hair cells at synaptic sites that are required for signaling to the brain. When nearby neurons release dopamine, activation of the dopamine receptors increases the activity of these mechanosensitive cells. A mutation in Aldh7a1 has been suggested to contribute in the Meniere disease (MD), an inner ear disorder characterized by tinnitus, vertigo, and hearing loss (Lynch et al., 2002). Acoustic overstimulation upregulate *Tyrp1* in rats (26520584). *Lbx1* acts as a selector gene in the fate determination of somatosensory and viscerosensory relay neurons in the hindbrain. Interestingly we found that Asip is upregulated in the The perception of sound involves the cochlear sensory epithelium (CSE), which contains hair cells and supporting cells. Hair cells are the transducers of auditory stimuli into neural signals, and are surrounded by supporting cells (PMID: 28662075). Taken together that these reports indicate that *Asip* a key regulator of several genes at different regions of the brain that play various role in developing and maintaining acoustic pathways.

### Sensory gating

#### Our results indicate a direct link between Hereditary hyperekplexia and aggression

Startle disease is characterized by an exaggerated startle response, evoked by tactile or auditory stimuli, leading to hypertonia and apnea episodes. Missense, nonsense, frameshift, splice site mutations, and large deletions in the human glycine receptor α1 subunit gene (GLRA1) are the major known cause of this disorder. However, mutations are also found in the genes encoding the glycine receptor β subunit (GLRB) and the presynaptic Na+/Cl−-dependent glycine transporter GlyT2 (SLC6A5). In this study, systematic DNA sequencing of SLC6A5 in 93 new unrelated human hyperekplexia patients revealed 20 sequence variants in 17 index cases presenting with homozygous or compound heterozygous recessive inheritance. Five apparently unrelated cases had the truncating mutation R439X. Genotype-phenotype analysis revealed a high rate of neonatal apneas and learning difficulties associated with SLC6A5 mutations. From the 20 SLC6A5 sequence variants, we investigated glycine uptake for 16 novel mutations, confirming that all were defective in glycine transport. Although the most common mechanism of disrupting GlyT2 function is protein truncation, new pathogenic mechanisms included splice site mutations and missense mutations affecting residues implicated in Cl− binding, conformational changes mediated by extracellular loop 4, and cation-π interactions. Detailed electrophysiology of mutation A275T revealed that this substitution results in a voltage-sensitive decrease in glycine transport caused by lower Na+ affinity. This study firmly establishes the combination of missense, nonsense, frameshift, and splice site mutations in the GlyT2 gene as the second major cause of startle disease.

LAMP5 LAMP5 plays a pivotal role in sensorimotor processing in the brainstem and spinal cord. It is highly expressed in several brainstem nuclei involved with auditory processing including the cochlear nuclei, the superior olivary complex, nuclei of the lateral lemniscus and grey matter in the spinal cord. It was localized exclusively in inhibitory synaptic terminals, as has been reported in the forebrain. Lamp5 knockout mice showed an increased startle response to auditory and tactile stimuli. In addition, LAMP5 deficiency led to a larger intensity-dependent increase of wave I, II and V peak amplitude of auditory brainstem response. (PMID: 30867010). We also found that Il33−/− animals had deficits in acoustic startle response, a sensorimotor reflex mediated by motor neurons in the brainstem and spinal cord (Fig. 2J, K; (23)). Auditory acuity and gross motor performance were normal (Fig. S5 I-J). Taken together, these data demonstrate that IL-33 is required for normal synapse numbers and circuit function in the thalamus and spinal cord (PMID: 29420261). Important (PMID: 25569367)

A number of controlled experiments and clinical investigations have demonstrated roles for glucocorticoids in auditory function and protection. As early as the 1960s, clinical studies revealed that patients with adrenocorticosteroid deficiency presented with greater auditory sensitivity compared to normal volunteers (Henkin et al., 1967). Moreover, treatment with prednisone brought hearing thresholds up to normal levels, demonstrating that the observed hypersensitivity was related to levels of circulating corticosteroids. Similarly other studies revealed that patients with Meniere’s disease, an inner ear disorder affecting both cochlear and vestibular function, exhibited low levels of circulating corticosteroids. Administration of adrenal cortex extract improved auditory function in these patients (Powers, 1972). One sphere in which steroids have an accepted and widespread use is in treatment of idiopathic sudden sensorineural hearing loss (Kuhn et al., 2011; Rauch et al., 2011). Exogenously administered synthetic glucocorticoids also protect the cochlea against damage induced by ototoxic drugs, acoustic trauma, and ischemia/reperfusion injury (Himeno et al., 2002; Takemura et al., 2004; Tabuchi et al., 2006). Given the transcriptional role of glucocorticoid receptors, several molecular changes likely underlie the observed protection. In particular, experiments point to enhanced biosynthesis of glutathione, reduced secretion of tumor necrosis factor induced cytokines, and altered expression of apoptotic genes as some of the changes likely to combat the free radical damage and apoptosis associated with noise- and chemically-induced cochlear damage (Maeda et al., 2005; Nagashima and Ogita, 2006; Hoang Dinh et al., 2009). (PMID:21909974)

## Acknowledgments

We would like to acknowledge *Professor Macrious Abraham* for his ideas and advice.

